# Molecular genetic analysis of Steroid Resistant Nephrotic Syndrome: Detection of a novel mutation

**DOI:** 10.1101/305987

**Authors:** Niloofar Serajpour, Behnaz Karimi, Nakisa Hooman, Rozita Hosseini, Pedram Khosravi, Hila Milo Rasouly, Azadeh Shojaei

## Abstract

Background: Nephrotic syndrome is one of the most common kidney diseases in childhood. About 20% of children are steroid-resistant NS (SRNS) which progress to end-stage renal disease (ESRD). More than 53 genes are associated with SRNS which represent the genetic heterogeneity of SRNS. This study was aimed to screen disease causing mutations within NPHS1 and NPHS2 and evaluate new potential variants in other genes.

Method: In first phase of study, 25 patients with SRNS were analyzed for NPHS1 (exon 2, 26) and all exons of NPHS2 genes by Sanger sequencing. In the second phase, whole exome sequencing was performed on 10 patients with no mutations in NPHS1 and NPHS2.

Result: WES analysis revealed a novel mutation in FAT1 (c.10570C>A; Q3524K). We identified 4 pathogenic mutations, located in exon 4 and 5 of NPHS2 gene in 20% of patients (V180M, P118L, R168C and Leu156Phe). Also our study has contributed to the descriptions of previously known pathogenic mutations across WT1 (R205C) and SMARCAL1 (R764Q) and a novel polymorphism in CRB2.

Conclusion: Our study concludes that mutations of exon 4 and 5 NPHS2 gene are common in Iranian and some other ethnic groups. We suggest conducting WES after NPHS2 screening and further comprehensive studies to identify the most common genes in the development of SRNS, which might help in Clinical impact on management in patients with SRNS.

**Detection of a novel mutation in SRNS**

## INTRODUCTION

Nephrotic Syndrome (NS) is one of the most common idiopathic primary diseases in childhood.(Bierzynska and Saleem 2017; Ha 2017; Weber et al. 2004) Which is defined as presence of four main symptoms: proteinuria, hyperlipidemia, hypoalbuminemia and edema (Bullich et al. 2015; Joshi et al. 2013). According to the patient’s response to the steroid therapy the disease divided into: resistant and sensitive groups. About 90% of patients are responsive to steroid therapy during four weeks who called steroid sensitive nephrotic syndrome (SSNS). Patients in whom proteinuria does not stop after about one month are classified as resistant which describe as steroid resistant nephrotic syndrome(SRNS)(Mekahli et al. 2009).SRNS is considered as a poor prognostic disease, in which 30-40% of it progresses to end stage renal disease(ESRD), requiring dialysis and transplantation(Tasic et al. 2015). The most frequent renal histological feature associated with SRNS is focal segmental glomerulosclerosis (FSGS). Moreover minimal change nephrotic syndrome (MCNS), and diffuse mesangial sclerosis (DMS) have been identified (Santin et al. 2009; Hinkes et al. 2007; Ha 2017). Genetic forms of SRNS are classified as isolated kidney disease or syndromic disorder (Ha 2017)The fenestrated endothelium, the glomerular basement membrane (GBM) and the podocytes form three layers of glomerular filtration barrier (GFB) which is impaired in NS and cause proteinuria.(Haraldsson et al. 2008)

2major proteins of podocytes including nephrin and podocin, coded by NPHS1 and NPHS2, are considered to play an important role in GFB. Mutations in these genes result in altering conformation and stability of podocytes and causing proteinuria and SRNS(Huber et al. 2003). Most cases of SRNS are considered as sporadic representing both AR and AD inheritance. NPHS1 and NPHS2 genes are the most common identified genes in AR form (Wang et al. 2017; Lovric et al. 2014).

This study was aimed to screen mutations causing disease within NPHS1 and NPHS2, figuring out the most common mutations in Iranian children and comprising the prevalence of such mutations among different nations. Due to heterogeneity of this disease, WES was performed for 10 patients in pilot study to evaluate other related genes and exploring new potential mutations. Indeed, preventing of ineffective treatment with steroids and helping proper clinician prediction in post transplantation outcome may be facilitated via indicating the specific mutations.

## 1. MATRIALS AND METODS

### 2.1 Patients’ description

25 subjects were recruited from Ali_Asghar children’s hospital in Tehran, Iran. The enrollment of patients in this study is based mainly on the clinical diagnosis of SRNS and disease onset ages varying from congenital to childhood. 3 of the children have been progressed into an end-stage renal disease and 5 of patients are undergoing dyalisis.The informed consent forms were given by parents of patients. The study was approved by the Ethical Committee of “Iran University of Medical Sciences, Faculty of Medicine”.

### 2.2 Polymerase chain reaction (PCR) and sequencing

4 ml of whole blood was taken from Patients and was transferred into tubes containing 200 μL EDTA for DNA isolation. DNA was isolated from peripheral blood of all samples by yekta tajhiz azma (YTA) kit (Iran).

All exons of NPHS2 gene were screened (primers are available upon request).exons 2 and 26 of NPHS1 gene were amplified using four primer pair (Supplementary data). Subsequently, to confirm the identified mutations in affected children, their parents were studied as well.

Reaction were accomplished in a total volume of 25 μL containing 12.5μl Master Mix (Amplicon with 1.5mM MgCl 2), 11 μl DEPC water, 20_40 ng template DNA and 10 pmol from each primer as well.

After initial denaturation at 94 °C for 3 min, 35 PCR cycles was performed using Thermofisher thermocycler thermocycler (SimpliAmp™ Thermal Cycler 96-well, Applied Bio systems); each cycle included denaturation at 94°C in 30s, annealing tempature at 62 for exon 2,26. extension at 72°C for 30s and final extension at 72°C in 5min.

PCR products were subjected to electrophoresis on agarose gel (1.5%).Sequencing was done by MACROGEN Company in South Korea using classic Sanger method with ABI. obtained sequences were aligned to the reference genome by chromas software and blast in Refseq in ncbi.

### 2.3 Whole exome sequencing (WES)

The second phase of study, WES was carried out for 10 patients by Colombia University Medical Center, IGM Institute for Genomic Medicine, Hemer Health Science.

## 2. RESULT

### 3.1 Polymerase chain reaction (PCR) and sequencing

25 children with SRNS were referred to Ali-Asghar hospital in Tehran, Iran to be examined for mutational analysis. Their mean age at the onset of symptoms was (2.54 ± 3.24) years (congenital to 14 years Positive family history was detected in 4 patients (16%), while 21 patients were sporadic(84%) in this cohort. Renal biopsy of patients indicated four different conditions, including FSGS (44%), MCNS (28%), CNF (8%), MeSPGN (4%).Histological data of 3 patients were not available. Also 9 patients showed positive family history of kidney stone. (Table.1)

**Table 1.** Clinical data of SRNS patients

| **Patient number** | **Gender** | **Sporadic/**<br>**Familial** | **Age at onset**<br>**(years/month)** | **Renal**<br>**biopsy** | **consanguinity** | **Family**<br>**history of**<br>**kidney**<br>**stone** |
| --- | --- | --- | --- | --- | --- | --- |
| **P1** | F | Sporadic | 3Y | MCNS | NO | NO |
| **P2** | F | familial | 2Y | FSGS | YES | YES |
| **P3** | M | sporadic | CONGENITAL | FSGS | NO | NO |
| **P4** | M | sporadic | 2Y | FSGS | YES | YES |
| **P5** | F | familial | 4Y | FSGS | NO | YES |
| **P6** | F | sporadic | 1.5Y | FSGS | YES | NO |
| **P7** | F | sporadic | 10Y | MeSPGN | YES | YES |
| **P8** | F | sporadic | 2Y | MCNS | YES | NO |
| **P9** | M | sporadic | 10M | FSGS | YES | YES |
| **P10** | F | familial | 7Y | MCNS | YES | YES |
| **P11** | M | sporadic | 3Y | – | YES | NO |
| **P12** | F | sporadic | 2Y | FSGS | YES | YES |
| **P13** | F | sporadic | 2Y | FSGS | NO | NO |
| **P14** | F | sporadic | 3Y | MCNS | YES | NO |
| **P15** | M | sporadic | 2.5Y | MCNS | NO | NO |
| **P16** | F | sporadic | CONGENITAL | CNS | YES | YES |
| **P17** | M | sporadic | 2Y | MCNS | YES | YES |
| **P18** | F | sporadic | 1Y | – | YES | YES |
| **P19** | F | sporadic | 5M | MCNS | YES | YES |
| **P20** | M | sporadic | CONGENITAL | – | YES | YES |
| **P21** | M | sporadic | 6M | FSGS | YES | YES |
| **P22** | M | sporadic | CONGENITAL | CNS | YES | NO |
| **P23** | F | familial | CONGENITAL | – | YES | YES |
| **P24** | F | sporadic | 8M | FSGS | YES | YES |
| **P25** | M | sporadic | 14.5Y | FSGS | NO | YES |

We identified c.567.568insT known pathogenic frameshift mutation (L156fsx166) in 2 patients. Moreover c.502C>T (p.168R>C) pathogenic homozygous mutation was found in one patient. Both of these mutations were located in exon 4 of NPHS2 gene. parents were screened for these mutations.(Figure 1) Although no mutation causing disease was detected in the other studied exons of NPHS1 and NPHS2 genes by this method, but benign or likely benign variants were detected in 15 patients (56%) within these regions.(Table.3)

**Fig 1.**
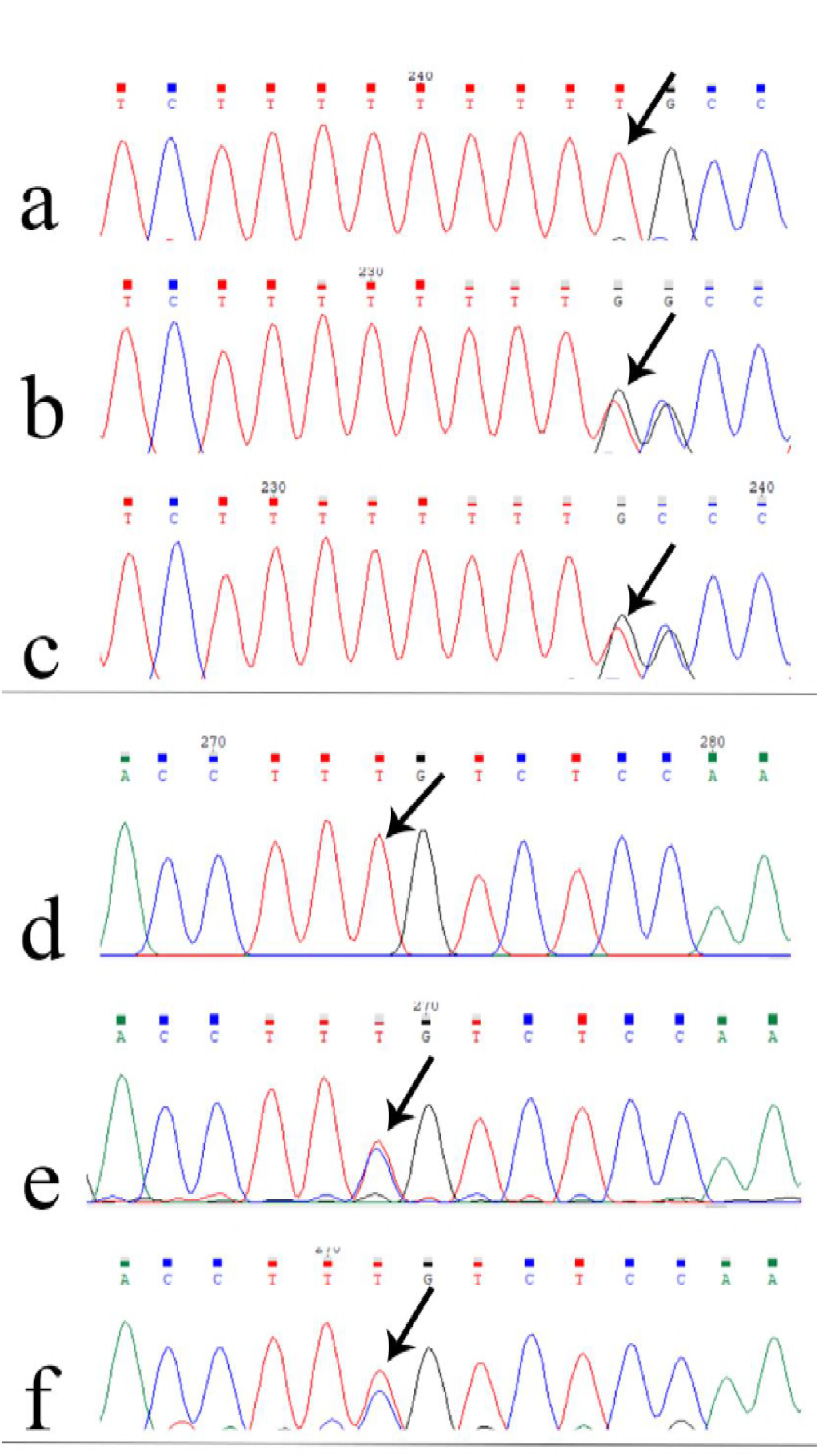
Found mutations in exon 4 of patients and parents. L156fsx166 (a, b and c) -p.168R>C (d, e and f)

### 3.2 Whole exome sequencing (WES)

To investigate other related genes and confer higher detection rate, WES was performed for 10 patients, with negative findings in the first phase of study. Two pathogenic mutations in NPHS2 were found in exon 2 and 5. Also, two other causative mutations were identified within WT1 and SMARCAL1 genes in 2 patients. Significantly, a novel mutation in FAT1 was detected. To predict the clinical significance of the found mutations, 3 different softwares were used, including SIFT, Polyphen, mutation assessor which revealed the prediction score of 0.034,0.044 and 2.1, respectively. (Table.2). Moreover, the parents of this patient were sequenced by PCR for confirmation of the novel mutation. (Figure 2) To amplify this region a specific primer pair was designed by primer3plus.(Supplementary data) Another novel variant were found in CRB2 predicted to be “ benign” by the above-mentioned silico analysis softwares.(0.058, 0.671,_) (Table.3).

**Table2.**
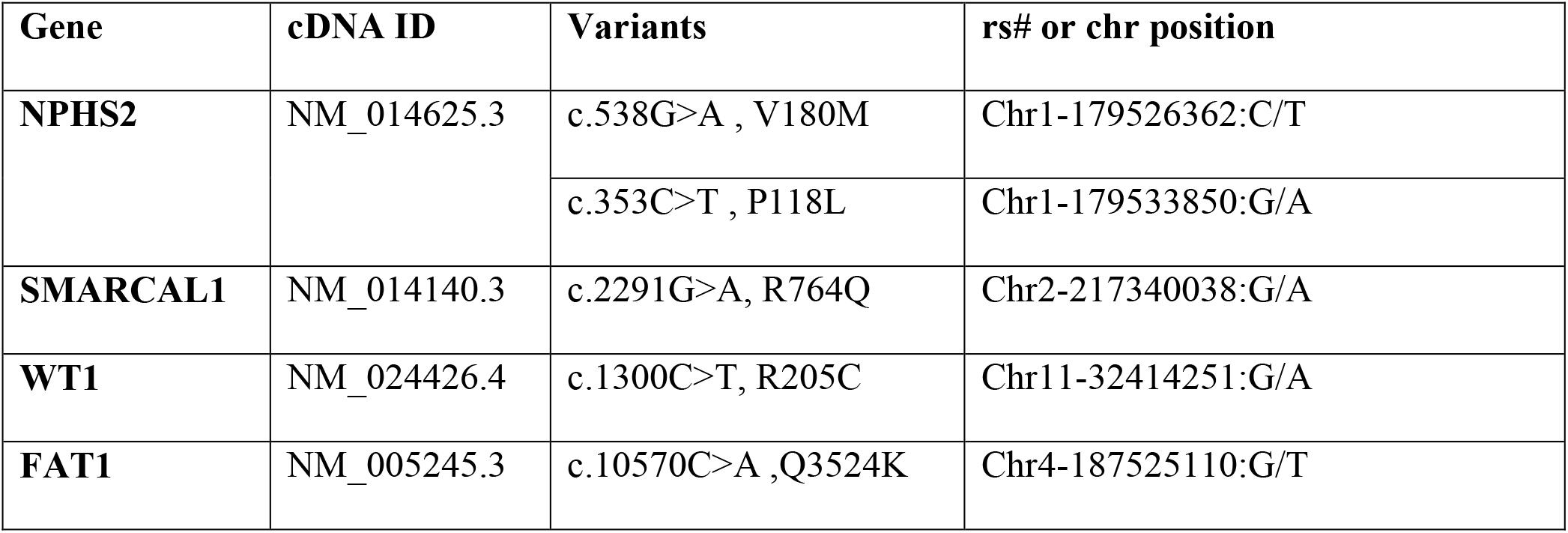
Found pathogenic mutations in the 5 patients with SRNS by WES

**Table3.** Found SNP in the 15 patients with SRNS

| Gene | Nucleotide change | Amino acid exchange | dbSNP name | Status (patient number) | References |
| --- | --- | --- | --- | --- | --- |
| **NPHS2** | c.954T > C | p.Ala318Ala | rs1410592 | Hom(1), Het(9) | (Karle et al. 2002) |
|  | c.1206C> G | 3UTR | rs1410591 | Hom(2), Het(6) | (Dusel et al. 2005) |
|  | IVS3 –46C > T | – | rs12401711 | Het(5) | (Aucella et al. 2005) |
|  | IVS3 –21C > T | – | rs12401708 | Het(5) | (Aucella et al. 2005) |
| **NPHS1** | c.3315G>A | p.Ser1105Ser | rs2071327 | Hom(1), Het(3) | (Shi et al. 2005) |
| **CRB2** | c.2259C>T | p.Ala753Val | - | Hom(1) | this study |

**Fig 2.**
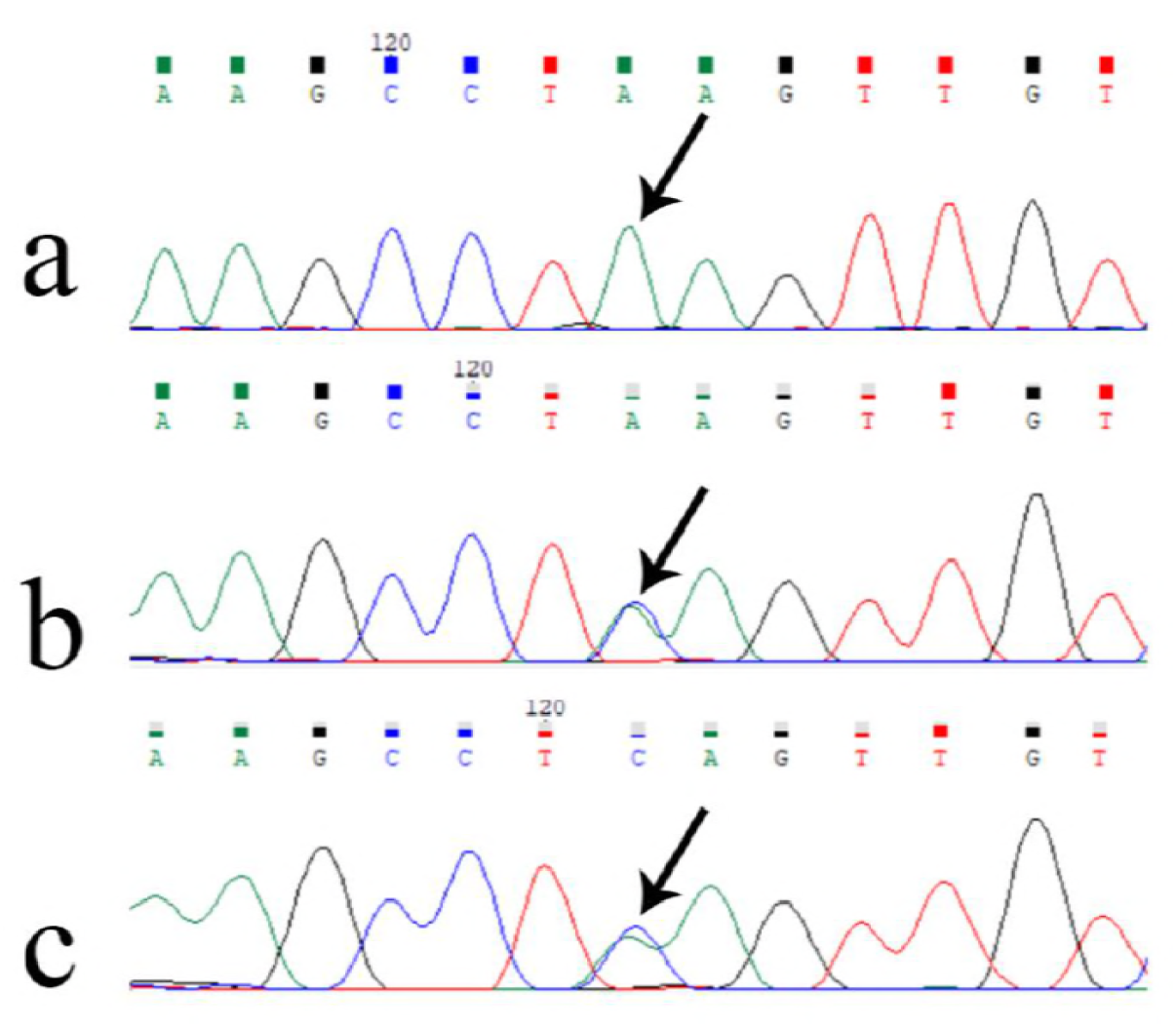
Novel mutation in FAT1 (a) and her parents (b and c)

## 3. DISCUSSION

Idiopathic nephrotic syndrome (INS) is a common clinical condition displaying genetic heterogeneity and significant phenotypic variability (Davin 2016; Bierzynska et al. 2017; Bullich et al. 2015; Sen et al. 2017). It can be caused by many single-gene mutations in both recessive (NPHS1, NPHS2, SMARCAL1, FAT1 and CRB2) and dominant inheritance forms (caused by genes such as WT1 and TRPC6).

Recessive mutations in NPHS1 and NPHS2 cause severe clinical features of early-onset NS and progress to ESRD, either during infancy or throughout childhood. Whereas, Hereditary autosomal-dominant NS is rare, occurring mostly in juvenile and adult familial cases(Ha 2017).

NPHS2 mutations have been reported as the most common cause of childhood onset in autosomal recessive SRNS (Sadowski et al. 2015; Guaragna et al. 2015; Tasic et al. 2015). The most frequent renal histological feature Associated with SRNS is focal segmental glomerulosclerosis

(FSGS)(Joshi et al. 2013; Liu and Wang 2017). In present study, the pathological manifestation of FSGS was about 44% in patients in which 4 out of 11patients with FSGS were showing NPHS2 mutation in a homozygous state.

Previous studies claim that incidence of NPHS2 mutations in children may vary according to ethnicity (Thomas et al. 2018; Guaragna et al. 2017). In 2013 Basiratnia et al (Basiratnia et al. 2013) showed that NPHS2 mutations were about 31%, responsible for 57% and 26% of sporadic and familial forms of SRNS. In our study, 5 of 25(20%) carried NPHS2 mutations (40% familial and 60% sporadic); both studies among Iranian population are almost consistent with findings of studies performed among American (Ruf et al. 2004a), Turkish (Berdeli et al. 2007), Arabic (Al-Hamed et al. 2013) and Mexican (Carrasco-Miranda et al. 2013) population. (26%, 24.7%, 22% and 21% respectively), while is in contrast to those findings among Far East population like Chinese (4.3%) (Yu et al. 2005), Japanese(4%) (Ogino et al. 2016) and south Korean children (0%) (Cho et al. 2008). due to all of these data, we suppose a hypothesis that the incidence of NPHS2 mutations decrease from northwest to southwest.

In present study we identified a known frameshift and a missense mutation in exon 4 of NPHS2 (Leu156Phe, R168C) in 3 patients (Karle et al. 2002; Weber et al. 2004). Previous investigations has been reported Two other mutations within this exon among Iranian population (R168H, D160G) (Ameli et al. 2012; Basiratnia et al. 2013). Although Otoukesh et al, indicated no mutation in exon 5 in 2009, but later 3 pathogenic mutations(V180M, R238S, F185fsX186) in this region were found by Basiratnia and her colleagues in 2013 (Basiratnia et al. 2013),similarly V180M in exon 5 was identified in one of our patient. Moreover, in our study P118L mutation was detected in exon 2. This missense mutation in podocin seems to be a relatively common NPHS2 mutation as it was found in several conducted studies.(Jaffer et al. 2014; Ruf et al. 2004a; Dincel et al. 2013) Although, Behnam et al reported there are more than 65% hot spot mutations in exon 8 of NPHS2, no mutation was found in our study within this region.

Despite the high rate of NPHS2 mutation, no hot spot mutations have been identified for this gene. But according to earlier (Basiratnia et al. 2013) and present study, we recommend NPHS2 especially exons4 and 5 should be considered as first step genetic approach in children with SRNS. Our finding is in consistent with among the other nations indicating common presence of SNPs within exon 4 and 5.(Berdeli et al. 2007; Karle et al. 2002; Tonna et al. 2008; Kari et al. 2013)

NPHS1 Mutations is another primary important gene associated with congenital nephrotic syndrome (CNS) that manifests within 90 days after birth with SRNS (Kestila et al. 1998). So far more than 200 mutations in NPHS1 have been reported (http://www.hgmd.org/, accessed on 2017). 2 known fin minor and fin major mutations in NPHS1 (within exon 26 and 2 respectively) have been found in majority of children (Yang et al. 2016). a study which was carried out in northwest of Iran by Behbahan et al (Behbahan et al. 2013) in 2013 reveled 6 different mutations in 80% of SRNS children showed no mutation within these exons. Similar to Brazilian (Guaragna et al. 2015) and polish (Binczak-Kuleta et al. 2014) studies, our result indicated pathogenic mutation neither within these exons among all patients nor in other exons of the cases studied by WES. Due to Behbahan’s findings (Behbahan et al. 2013) and our study, we suppose that exon 2 and 26 of NPHS1 gene may not be as causative exons for SRNS in Iranian children.

It is acknowledged that more than 53 genes are associated with SRNS in both recessive and dominant inheritance form (Bierzynska et al. 2014; Sen et al. 2017). Gemma Bullich et al supposed that genetic testing using standard Sanger methods is costly and time consuming, even if only the most frequently mutated genes are analyzed (Bullich et al. 2015) but we think that screening for pathogenic variants in some common genes by this method could be the first cost effective approach. Due to genetic heterogeneity of SRNS, Next step may be employing WES. It is noted although WES cost about 30% more but it leads to identification of new disease-causing mutations covering all genes associated with SRNS (Wang et al. 2017).

In our study, through evaluation of WES data in patient 16, known R764Q mutation was found in SMARCAL1 gene, a transcription factor express in podocyte(Antignac 2005). This mutation is related to a disorder known as Schimke immune-osseous dysplasia (SIOD) showing SRNS, short stature and immunedeficiency (Lowik et al. 2009; Boerkoel et al. 2002). Finding a putative mutation using WES method in this case helped us diagnose a disease which its symptoms overlap with SRNS.

Another gene involved in SRNS is FAT1 which Loss of function mutations in this gene results in decreased cell adhesion and migration in fibroblasts and podocytes (Gee et al. 2016). Most of previous studies describe the role of FAT1 heterozygous mutations in some cancers (Morris et al. 2013; Comprehensive genomic characterization of head and neck squamous cell carcinomas 2015), whereas Heon Yung Gee and collogues in 2016 reported 4 different homozygous variant as a causative factor for glomerulotubular nephropathies such as NS (Gee et al. 2016). We here identified a novel recessive variant Q3524K in FAT1 as causing SRNS in a 10 years girl from consanguine parent. This potentially pathogenic variant was evaluated by some predictive tools including SIFT, Polyphen and Mutation Assessor. However to confirm its pathogenicity some functional and in vitro investigations are needed.

Mutations in the WT1 gene, encoding the Wilms’ tumor 1 protein, which typically lead to Denys-Drash syndrome or Frasier syndrome, can also cause SRNS Type 4 (Chernin et al. 2010; Miller-Hodges 2016; Hall et al. 2015).

Exon 8 and 9 of this gene has been considered as one the most prominent implicated genes in SRNS (Mucha et al. 2006; Sen et al. 2017). Clinical manifestation of patient 18 showed an affected girl with congenital and sporadic CNS/SRNS. In this index, WES analysis revealed one homozygous R205C mutation in exon 7, which was a de novo mutation in a hot spot region (Lipska et al. 2014). Our findings are in consistent with previous reports identifying WT1 mutation mostly in girls, within hot spot regions of WT1 gene (exons 5, 6, 7, 8 and 9) and often in de novo state (Kumar et al. 2016; Lowik et al. 2009).

The low rate of mutation frequency in WT1 gene of our study (4%) is similar to some reports by Ruf and his collogues(Ruf et al. 2004b), cho et al (Cho et al. 2008) and Alharthi et al (Alharthi et al. 2017) (6.9%, 5.7% and 5% respectively)but is less than reports of MUCKA study(8.9%) (Mucha et al. 2006) and more than the finding related to Indian children(1.7%). Although, mutation rate of WT1 is low but patients carrying WT1 mutations represent in early onset with more severe phenotype and congenital form of SRNS.

In overall, SRNS cause 15% of all chronic kidney disease. In our study, we noticed the interesting data that positive family history of Kidney stone were existed in 16 out of 25(64%) patients. This finding triggered a hypothesis in our mind that this factor may increase the risk of NS significantly, although to prove its accuracy, more samples and further investigations should be performed.

## 4. Conclusion

In summary, this is the first and largest study among Iranian population with different ethnic origins that investigates causative variants associated with SRNS through screening both common genes (NPHS1 and NPHS2) and whole exome study. Among 25 patients who underwent for PCR sequencing for all exons of NPHS2, 5 patients carried a mutation causing disease, suggesting that NPHS2 especially exons 4 and 5 of this gene should be considered as the first step genetic approach in children with SRNS. For the first time in this country, 3 known variant were detected in WT1, SMARCAL1 and CRB2, significantly a novel variant were identified in FAT1 gene.

Because of the heterogeneous clinical and pathological spectrum, a molecular diagnosis based on sequencing is required. Identification of mutations causing SRNS is of importance, not only for therapeutic considerations but also for genetic counseling.

## ACKNOWLEDGMENTS

This study was approved and supported by grant no. 96-03-208-31481 by Research Deputy, IUMS. We gratefully thank Dr.Ali Gharavi for conducting WES and his technical support (Colombia University Medical Center, IGM Institute for Genomic Medicine, Hemer Health Science).

